# Norepinephrine is a novel and essential regulator for T-lymphocyte interleukin 17A expression

**DOI:** 10.64898/2026.08.20.746026

**Authors:** Tamara Natour, Tatlock H. Lauten, Lauren J. Pitts, Emily C. Reed, Kristen R. Giebel, Adam J. Case

## Abstract

The longstanding association between psychological trauma or stress and subsequent chronic inflammatory disorders is well-documented, but the mechanism by which psychopathology causes these immune changes has yet to be elucidated. We have previously reported that sympathetic innervation to lymphoid organs and beta adrenergic receptor signaling are essential for T-lymphocyte interleukin 17A (IL-17A) production and T_H_17 polarization, though the exact mechanistic contribution of this signaling to the development of T_H_17 cells remained unclear. Therefore, we hypothesized that norepinephrine (NE) is a novel and direct regulator of T-lymphocyte IL-17A expression. Herein, we indeed observed that NE regulates baseline IL-17A *in vivo*. We further identified a novel mechanism by which transforming growth factor beta (TGFβ) and NE together result in T_H_17 polarization and IL-17A production in CD4+ T-lymphocytes. Additionally, we found that cAMP, PKA, and CREB are induced by NE signaling, which ultimately increases CBP/p300 activity to initiate RORγt transcription. Surprisingly, our data also demonstrate that STAT3, a transcription factor previously described as necessary for canonical T_H_17 polarization, is not involved in this novel pathway and may even be downregulated. Combined with bulk RNA sequencing data, our data highlight robust differences between the novel and canonical pathways to T_H_17 polarization. Altogether, our data reveal a novel, noncanonical mechanism that links sympathetic nervous system activity with IL-17A related inflammation, which may have significant relevance to sympathoexcitation-related disorders that stem from psychological trauma or stress.

## Introduction

The crosstalk between psychological trauma and immune system dysregulation is slowly becoming more appreciated. While it is known that individuals diagnosed with post-traumatic stress disorder (PTSD), a potential consequence of psychological trauma, are more likely to develop subsequent chronic inflammatory conditions (Boscarino et al., 2010; Passos et al., 2015; Speer et al., 2018; Wang et al., 2017), the precise causal signaling mechanisms remain poorly understood. Interleukin 17A (IL-17A) is a proinflammatory cytokine that is intimately linked to many autoimmune and inflammatory pathologies (Majumder and McGeachy, 2021; Mills, 2023) and is also elevated in humans following psychological trauma (Sommershof et al., 2009; Zhou et al., 2014; Zuo et al., 2023). Its primary producer is the T_H_17 T-lymphocyte, and prevailing evidence suggests canonical T_H_17 polarization requires naïve CD4+ T-lymphocytes to activate in the presence of TGFβ, IL-1β, IL-23, and IL-6 (Hanna et al., 2022), with particular emphasis placed on the necessity of TGFβ and IL-6 (Heink et al., 2017; Zhou et al., 2007). This cytokine milieu is generally produced in response to specific pathogens where it is believed that downstream production of IL-17A from T-lymphocytes plays a supportive role in coordinating both innate and adaptive immunity (Khader et al., 2009; Lyakh et al., 2008; Zielinski et al., 2012). However, while psychological trauma induces IL-17A, it is not reported to upregulate all the aforementioned canonical cytokines needed for its expression, which suggests there may be additional unknown signals able to generate IL-17A from T-lymphocytes.

Using a psychological trauma mouse model known as repeated social defeat stress (RSDS), we have previously recapitulated the human phenomenon of significant elevations in circulating IL-17A and T_H_17 T-lymphocytes (Elkhatib et al., 2020; Moshfegh et al., 2019). Elevated sympathoexcitation and norepinephrine (NE) are known signatures of psychological trauma and PTSD (Bremner and Wittbrodt, 2020; Geracioti et al., 2001; Kosten et al., 1987), and we previously demonstrated that both sympathetic denervation and eliminating the ability of T-lymphocytes to produce catecholamines significantly blunted T-lymphocyte IL-17A production after RSDS (Elkhatib et al., 2022; Elkhatib et al., 2021). Furthermore, we have shown that both pharmacological and genetic inhibition of beta 1- and 2-(β1/2) adrenergic receptors on T-lymphocytes completely prevented T_H_17 polarization and IL-17A production both at baseline and following RSDS (Lauten et al., 2024). Together, these data suggest that a sympathetic-mediated ligand working through β-adrenergic signaling on T-lymphocytes is critical for IL-17A expression, and we hypothesize this ligand to be NE.

Herein, we investigated this hypothesis by exploring the necessity and sufficiency of NE on T_H_17 polarization. We found a direct correlation between systemic NE and IL-17A levels, with depletion of NE leading to an equivalent loss of IL-17A *in vivo*. While NE alone was not sufficient to generate IL-17A from T-lymphocytes, it was able to induce IL-17A in combination with transforming growth factor-beta (TGFβ), though not to the magnitude of the canonical T_H_17 polarization cytokine cocktail. This novel TGFβ/NE induction of IL-17A appears to signal via cyclic AMP (cAMP), protein kinase A (PKA), and ultimately cAMP response element-binding protein (CREB), with the signals converging at the transcriptional coactivators CBP and p300 (Janknecht et al., 1998; Pouponnot et al., 1998). Interestingly, by RNA sequencing, we elucidated that TGFβ/NE and canonical T_H_17 polarization led to increased IL-17A production from T-lymphocytes by different signaling pathways. Altogether, these findings suggest a synergistic effect leading to IL-17A produced by the convergence of NE and TGFβ signaling, which may also contribute to the chronic inflammation due to psychological trauma and enhanced sympathoexcitation.

## Materials and Methods

### Mice

Wild-type C57BL/6J (#000664; shorthand WT) and IL-6 knock-out (#002650; shorthand IL6^-/-^) mice were obtained from Jackson Laboratories (Bar Harbor, ME, USA). All mice were bred in house to eliminate shipping stress and microbiome shifts, as well as co-housed with their littermates (≤5 mice per cage) prior to the start of experimentation to eliminate social isolation stress. Mice were housed with standard pine chip bedding, paper nesting material, and given access to standard chow (#8604 Teklad rodent diet, Inotiv, West Lafayette, IN, USA) and water ad libitum. Male and female experimental mice between the ages of 8-12 weeks were utilized in all experiments, but no sex differences were observed so data are presented as pooled independent of sex. Experimental mice were randomized, and when possible, experimenters were blinded to the respective cohorts until the completion of the study. Mice were sacrificed by pentobarbital overdose (150mg/kg, Fatal Plus, Vortech Pharmaceuticals, Dearborn, MI, USA) administered intraperitoneally. All mice were sacrificed between 7:00 and 9:00 Central Time to eliminate circadian rhythm effects on T-lymphocyte function. All procedures were reviewed and approved by Texas A&M University Institutional Animal Care and Use Committees.

### In vivo experimentation

6-hydroxydopamine (#HY-B1081A, MedChem Express, Monmouth Junction, NJ, USA) was reconstituted with 0.1% ascorbic acid in 1X PBS and delivered via two intraperitoneal injections administered on days 0 and 2 using the previously established dose of 100 mg/kg (Gomez-Flores et al., 2021).

### T-lymphocyte immunophenotyping

Flow cytometric immunophenotyping was performed as previously described(Lauten et al., 2024). Briefly, cultured CD4+ T-lymphocytes were harvested and incubated at 37 °C for 4 h in RPMI media supplemented with phorbol 12-myristate-13-acetate (PMA; 10 ng/mL), ionomycin (0.5 mg/mL), and BD GolgiPlug Protein Transport Inhibitor (containing brefeldin A; 1 mg/mL; BD Biosciences, Franklin Lakes, NJ, USA). Following this, cells were washed and resuspended in PBS containing an amine-reactive viability dye (Live/Dead Fixable Cell Stain, #L34960, Thermo Fisher Scientific, Houston, TX, USA) for 30 min at 4 °C. An antibody targeting intracellular RORγt (#5016098, ThermoFisher Scientific) was then added following fixation/permeabilization (#00-5523-00, eBioscience, San Diego, CA, USA). Cells were assessed using a 4-laser Attune NxT flow cytometer (Thermo Fisher Scientific, Houston, TX, USA), while data was analyzed using FlowJo analysis software (BD Biosciences, Franklin Lakes, NJ, USA).

### T-lymphocyte isolation and activation

T-lymphocytes from spleens were isolated using negative selection as previously described (Moshfegh et al., 2019). Briefly, splenocytes in a single cell suspension were negatively selected for naïve CD4+ T-lymphocytes using the EasySep Mouse Naïve CD4+ T-cell Isolation Kit (StemCell Technologies #19765, Vancouver, BC, USA). Cell viability was measured via a Bio-Rad TC20 Automated Cell Counter using trypan blue exclusion, and cell purity was assessed via flow cytometry (average yield is approximately 1 × 10^6 cells/mouse and purity achieved is >90 %). T-lymphocytes were cultured in RPMI media supplemented with 10 % Fetal Bovine Serum, 10 mM HEPES, 2 mM Glutamax, 100 U/mL penicillin/streptomycin, and 50 μM of 2-mercaptoethanol.

### Canonical T_H_17 polarization

CD4+ splenic T-lymphocytes were polarized to IL-17A-producing phenotypes *ex vivo* by use of CellXVivo Mouse T_H_17 Cell Differentiation Kit (#CDK017, R&D Systems, Minneapolis, MN, USA) as previously described(Lauten et al., 2024). Briefly, CD4+ T-lymphocytes were isolated and cultured as described above with RPMI media supplemented with Dynabeads Mouse T-Activator CD3/CD28 (Gibco #11452D) at a ratio of 1:1 cell to beads, mouse IL-6 (#130-096-682, Miltenyi Biotech, Bergisch Gladbach, North Rhine-Westphalia, Germany), mouse IL-23 (#130-096-676, Miltenyi Biotech, Bergisch Gladbach, North Rhine-Westphalia, Germany), mouse IL-1β (#130-101-681, Miltenyi Biotech, Bergisch Gladbach, North Rhine-Westphalia, Germany), human TGF-β1 (#130-095-067, Miltenyi Biotech, Bergisch Gladbach, North Rhine-Westphalia, Germany), anti-mouse IL-4 antibody (#130-095-709, Miltenyi Biotech, Bergisch Gladbach, North Rhine-Westphalia, Germany) anti-mouse IFN-γ antibody (#130-095-729, Miltenyi Biotech, Bergisch Gladbach, North Rhine-Westphalia, Germany), and anti-mouse IL-2 antibody (#130-095-736, Miltenyi Biotech, Bergisch Gladbach, North Rhine-Westphalia, Germany). Following varying time points of culture and activation, T-lymphocytes were harvested for analysis.

### In vitro treatment regiments

Cultured T-lymphocytes were treated with L-Norepinephrine (#AAL0808703, Thermo Scientific Chemicals, Houston, TX, USA) and human TGF-β1 (#130-095-067, Miltenyi Biotech, Bergisch Gladbach, North Rhine-Westphalia, Germany) to induce noncanonical T_H_17 polarization. Cells were also supplemented with A-485 (#HY-107455, MedChem Express, Monmouth Junction, NJ, USA), MDL-12330A (#444200, Sigma Aldrich, St. Louis, MO, USA), H89 (#B1427, Sigma Aldrich, St. Louis, MO, USA), and/or Forskolin (#F3917, Sigma Aldrich, St. Louis, MO, USA). Concentrations used are denoted in specific figures and were based on previously established doses (Lauten et al., 2024).

### Catecholamine quantification

Catecholamines were quantified as previously described(Lauten et al., 2024). Briefly, dopamine, norepinephrine, and epinephrine levels in splenic lysates and plasma were assessed using the 3-CAT research ELISA (Rocky Mountain Diagnostics, BAE-5600, Colorado Springs, CO, USA). All assays were completed according to the manufacturer’s protocol.

### Chromatin Immunoprecipitation and ChIP-qPCR

Chromatin immunoprecipitation was performed using the low cell ChIP kit (#53086, Active Motif, Carlsbad, CA, USA). Assays were completed according to the manufacturer’s protocol. RT-qPCR was performed to assess DNA associated with the RORC promoter, CNS6, and CNS9 loci. Primers for these loci were derived from Chang et al (Chang et al., 2020). DNA enrichment was reported as fold change of percent of input. Primer sequences listed below:

Rorc P; Forward: GGGGAGAGCTTTGTGCAGAT; Reverse: AGTAGGGTAGCCCAGGACAG

CNS6; Forward: GCCCACCAATCTGTCCCCACTA; Reverse: TTCTGACTCGCTTTCCTCCCAG

CNS9; Forward: GCACAGGACCAGAACAAGCAGG; Reverse: AGCACAAGAAACGGGGAGACTGA

### cAMP quantification

As previously described (Lauten et al., 2024), quantification of cAMP levels within cell lysates was assessed using the cAMP research ELISA (#ab290713, Cambridge, United Kingdom). All assays were completed according to the manufacturer’s protocol, with cell lysate normalized to live cell counts.

### RNA extraction, cDNA production, and quantitative real-time RT-PCR

T-lymphocyte gene expression was quantified as previously described(Moshfegh et al., 2023). Briefly, mRNA was extracted using a RNAeasy plus mini kit (#74136, Qiagen) and quantified using NanoDrop One Spectrophotometer (#13400518, Thermo Scientific). RNA was then transformed into cDNA using ThermoFisher High-Capacity cDNA Reverse Transcriptase Kit (#4374967, Applied BioSystem). Generated cDNA was used for real time quantitative PCR. Primers for 18s and IL-17A were designed using NIH primer-BLAST spanning exon-exon junctions. Cq values were determined, and relative gene expression was calculated by comparing housekeeping 18s ribosomal gene expression to gene of interest (2^-ddCq). Primer sequences listed below:

18s; Forward: GCCCGAAGCGTTTACTTTGA; Reverse: TCATGGCCTCAGTTCCGAA

Il17a; Forward: CAGGGAGAGCTTCATCTGTGT; Reverse: GCTGAGCTTTGAGGGATGAT

### Protein analysis

Protein was isolated and assessed as previously described(Reed et al., 2024). Briefly, T-lymphocyte protein was isolated using RIPA lysis buffer (#PI89900, Thermo Scientific) and 1% Halt protease inhibitor cocktail (#PI87785, Thermo Scientific). Samples were subsequently subjected to sonication and centrifugation to obtain soluble protein, which was quantified using Pierce BCA protein assay kit (#PI23227, Thermo Scientific). Mouse CREB and STAT3 protein was assessed via Total Protein Jess Automated Western Blot (Bio-techne) as previously described(Moshfegh et al., 2023), using Rabbit anti-CREB, anti-pCREB (Ser133), anti-STAT3, and anti-pSTAT3 (Tyr705) primary antibodies (#4820S #9198S, #12640S, #9145S, Cell Signaling, respectively). Total protein and phospho-protein antibodies were diluted 1:100 and 1:50, respectively. Analysis was performed using Compass Software for Simple Western.

### Cytokine analysis

Cytokines in plasma and media (normalized to cell counts) were measured as previously described(Elkhatib et al., 2020). Briefly, concentrations were assessed using custom Mesoscale Discovery U-Plex kits (Mesoscale Discovery, Rockville, MD, USA) per manufacturer’s instructions. Analysis was performed using a Mesoscale QuickPlex SQ 120 and analyzed using Mesoscale Discovery software.

### Bulk RNA sequencing

Bulk RNA sequencing was performed as previously described (Reed et al., 2025). Briefly, Splenic CD4+ T-lymphocytes were isolated and activated with Dynabeads for 24 hours. RNA was extracted using the RNAeasy plus mini kit (#74136, Qiagen) with DNAse treatment (#79254, Qiagen) and quantified using NanoDrop One Spectrophotometer (#13400518, Thermo Scientific). The eight RNA samples were assessed for quality using the TapeStation system, and RNA concentrations were measured using the Qubit fluorometer. All samples had an RNA Integrity Number (RIN) greater than 9. Approximately 180 ng of total RNA from each sample was used for rRNA-depleted RNA-seq library preparation with the WATCHMAKER RNA Library Prep Kit with Polaris Depletion, following the manufacturer’s protocol. The libraries were amplified using 10 cycles of PCR and subsequently evaluated on the TapeStation system, yielding an average library size of approximately 295 bp. The pooled libraries were sequenced on an Illumina NextSeq 2000 using a P2 200-cycle flow cell with a paired-end 2 × 100 bp sequencing configuration. Raw reads were trimmed and assessed using Trim-Galore! (v0.6.10) and MultiQC (v1.30). Quality metrics indicated adequate read quality for downstream analyses. Reads were aligned to the mouse reference genome (GRCm39) with STAR (v2.7.11b), and gene level counts were obtained using featureCount from the Subread package (v2.0.8). The resulting count matrix was processed in DESeq2 (v1.48.1) for differential expression analysis. Comprehensive pathway enrichment was performed using Gene Ontology (GO), KEGG, Reactome, and Gene Set Enrichment Analysis (GSEA). Raw sequencing data is available in the Texas Data Repository: https://dataverse.tdl.org/dataset.xhtml?persistentId=doi:10.18738/T8/R6W6IA.

## Statistical analysis

No sex differences in parameters measured were observed, so data are pooled independent of sex. All data are presented as mean ± standard error of the mean (SEM) with sample numbers displayed as individual data points and N values are included within figure legends where appropriate. Data were first assessed for normality utilizing the Shapiro-Wilk test. Following this, data were analyzed using Mann-Whitney U test, 1-way ANOVA, and 2-way ANOVA where appropriate (specific tests identified in the figure legend). All statistics were calculated using GraphPad Prism (V11, GraphPad).

## Results

### Norepinephrine is a regulator of IL-17A production in the presence of TGFβ

IL-6 is a critical cytokine in canonical T_H_17 polarization (Bettelli et al., 2006; Veldhoen et al., 2006). Yet, we observed IL-6 knock-out (IL6^-/-^) mice maintain basal levels of IL-17A. To investigate the role of NE in IL-17A expression, we depleted peripheral NE levels by 6-OHDA mediated chemical sympathectomy. Strikingly, basal levels of plasma IL-17A were significantly attenuated with 6-OHDA administration in both wild-type (WT) and IL-6^-/-^ animals (**Fig. 1A**). NE was confirmed to be depleted with 6-OHDA administration in both the plasma and spleens of animals (**Supplementary Fig. 1A-B**), which also revealed a positive correlation between plasma IL-17A and NE levels (**Supplementary Fig. 1C**). To explore how NE drives IL-17A production, naïve CD4+ T-lymphocytes were activated *in vitro* and supplemented with each individual cytokine with or without NE. Of note, NE alone was not sufficient to increase IL-17A expression significantly over control (**Fig. 1B**). Interestingly, TGFβ was the only cytokine that, in combination with NE, produced a significant increase in media IL-17A (**Fig. 1B**). It was observed that 2 ng/mL TGFβ + 10 µM NE provided the most robust and reproducible IL-17A expression (**Fig. 1C-D**), therefore, this dose was used throughout the remaining studies and will be referred to as the noncanonical T_H_17 pathway. To assess the necessity of autocrine/paracrine IL-6 on T-lymphocyte IL-17A expression, IL6^-/-^ naïve CD4+ T-lymphocytes were polarized via the noncanonical T_H_17 pathway. Though not to the same magnitude as WT T-lymphocytes, IL6^-/-^ T-lymphocytes were able to produce IL-17A under the noncanonical conditions (**Fig. 1E**), suggesting the involvement of a separate (though potentially additive) mechanism of IL-17A induction that is independent of IL-6 signaling. Additionally, the noncanonical pathway induces significantly less IL-17A production compared to the canonical T_H_17 conditions (**Fig. 1F**). This suggests that the canonical cytokine milieu produces a much larger and more robust IL-17A response, while the noncanonical pathway may be more consistent with low-grade inflammation.

**Figure 1:**
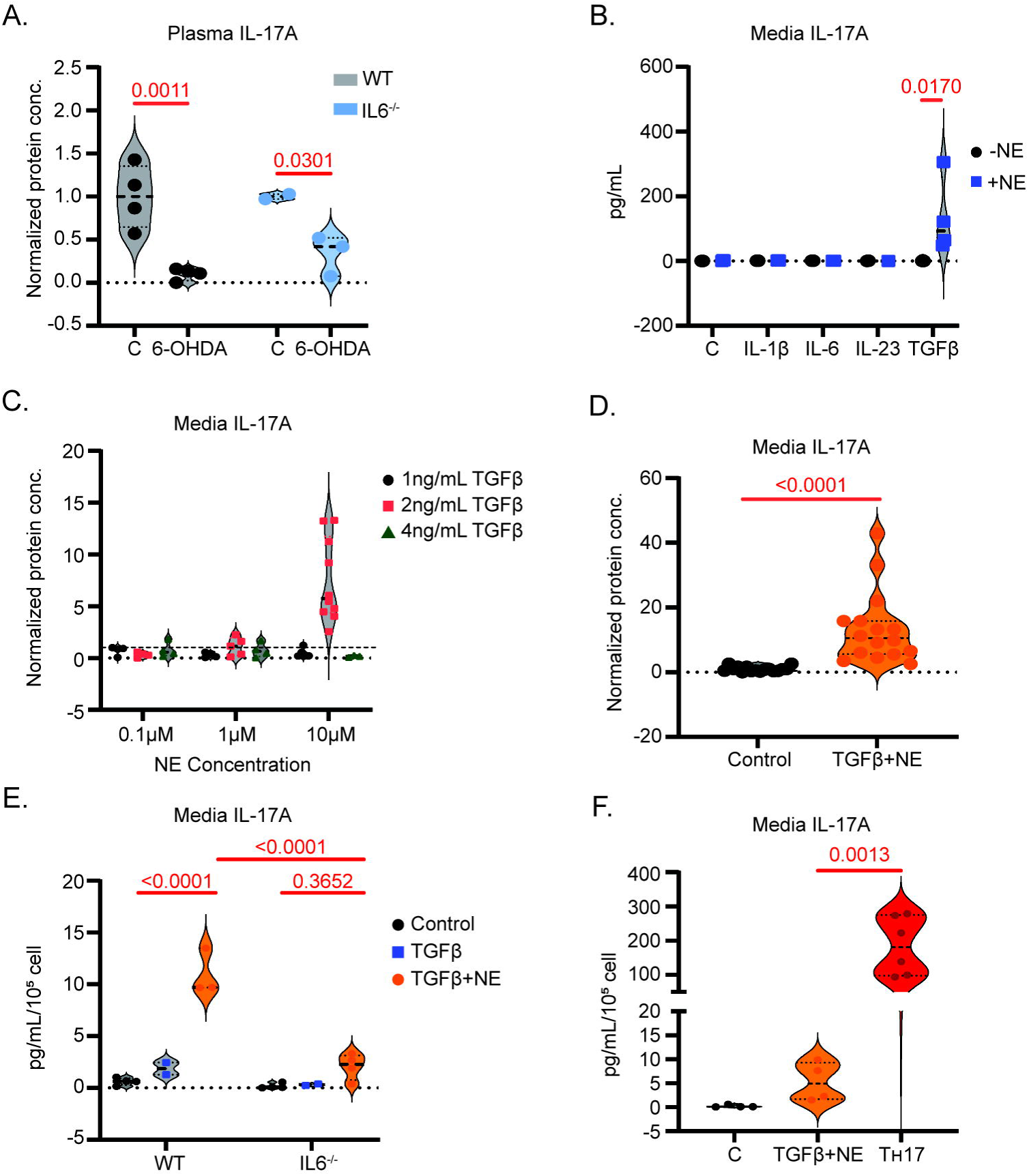
NE is essential for in vivo basal IL-17A levels and is sufficient for in vitro IL-17A production with TGFβ. Plasma IL-17A protein in WT (N=8) or IL6^-/-^ (N=5) mice one-week post-6-OHDA injections. Data normalized to control animals within genotype **(A)**. Media IL-17A protein 72 hours after activation and treatment of naïve CD4+ T-lymphocytes with respective treatments per panel **(B-F)**. Statistics were measured by 2-way ANOVA with Tukey’s multiple comparisons test (A,C,E), unpaired t-test (D), or ordinary one-way ANOVA with Tukey’s multiple comparisons test (B,F).

### cAMP, PKA, and CREB, but not STAT3, are involved in the noncanonical polarization pathway

Due to the established role of G-protein coupled receptor (GPCR) activation of adenylyl cyclase in adrenergic signaling, we first explored noncanonical T_H_17 intracellular signaling by assessing intracellular cyclic AMP (cAMP) levels in T-lymphocytes under the canonical and noncanonical T_H_17 conditions. As expected with NE stimulation, the noncanonical pathway exhibited significantly higher levels of cAMP compared to control and canonical T_H_17 conditions (**Fig. 2A**). To confirm that cAMP is a critical component of noncanonical IL-17A production, naïve T-lymphocytes were supplemented with TGFβ and forskolin, which yielded similar increases in IL-17A mRNA to TGFβ and NE (**Fig. 2B**). This was further supported by the complete prevention of IL-17A production by the adenylyl cyclase and downstream protein kinase A (PKA) inhibitors, MDL-12330A and H89, respectively (**Fig. 2C**). These data warranted the measurement of cAMP response element-binding protein (CREB) activity under the noncanonical conditions. Once again, the noncanonical pathway significantly elevated CREB activity compared to canonical conditions, as evidenced by a higher phospho-CREB to total CREB ratio (**Fig. 2D-E**).

**Figure 2:**
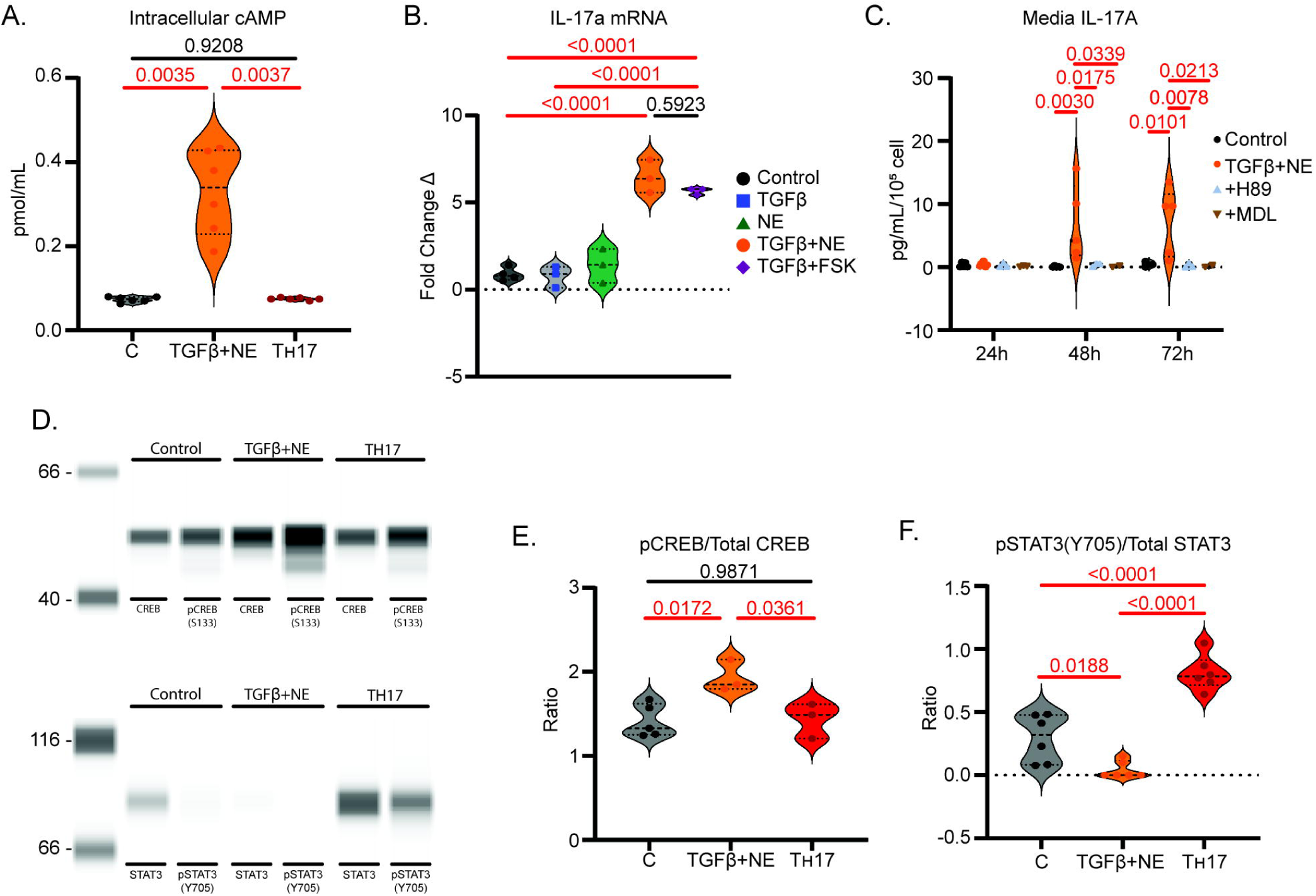
cAMP, PKA, and CREB, but not STAT3, are involved in the noncanonical T_H_17 pathway. Intracellular cAMP levels 15 minutes after treating naïve CD4+ T-lymphocytes with TGFβ+NE (N=6) or canonical T_H_17 cocktail (N=6) **(A)**. IL17a mRNA 72 hours after treating naïve CD4+ T-lymphocytes with TGFβ, NE, both, or TGFβ and 1 µM FSK (N=3 per group) **(B)**. Media IL-17A protein normalized to cell counts 24-, 48-, and 72-hours after treating naïve CD4+ T-lymphocytes with TGFβ+NE alone, with 10 µM H89 (PKA inhibitor), or with 10 µM MDL-12330A (adenylyl cyclase inhibitor) **(C)**. 30-minute SimpleWestern Automated Jess of pCREB/total CREB protein measurements in control (N=5), TGFβ+NE treated (N=3), or canonical T_H_17 treated (N=3) CD4+ T-lymphocytes **(D, E).** 72-hour SimpleWestern Automated Jess pSTAT3/total STAT3 protein measurements in control (N=6), TGFβ+NE treated (N=6), or canonical T_H_17 treated (N=6) CD4+ T-lymphocytes **(D, F)**. Statistics were measured by one-way ANOVA with Tukey’s multiple comparisons test (A,B,E,F) or 2-way ANOVA with Tukey’s multiple comparisons test (C).

Classically, the transcription factor STAT3 is thought of as critical and indispensable for T_H_17 polarization by way of canonical signaling (Bhaumik and Basu, 2017; Dong, 2008; Yang et al., 2007; Zhou et al., 2007). Therefore, we interrogated the activation of STAT3 via phosphorylation at Y705 (pSTAT3) in the noncanonical system. Astonishingly, not only was pSTAT3 absent, but total STAT3 was also reduced with TGFβ and NE treatment compared to control (**Fig. 2D, F**), further suggesting diverging mechanisms between canonical and noncanonical T_H_17 polarization.

### TGFβ+NE converge at CBP/p300 and synergistically increase RORγt transcription

TGFβ is known to signal through a well-established pathway involving Smad proteins, in which Smad2 and 3 are phosphorylated and associate with Smad4 (Shi and Massagué, 2003). This complex then translocates into the nucleus where it regulates transcription of RORγt and IL-17A (Shi and Massagué, 2003; Zhang, 2018). It has also been reported that TGFβ-induced Smad transcriptional activity is enhanced through the recruitment of the coactivators CBP and p300 (Janknecht et al., 1998; Pouponnot et al., 1998; Shen et al., 1998), which has been shown to target regulatory regions at the Rorc locus and act as a histone acetyltransferase (HAT) allowing for transcription of RORγt (Yahia-Cherbal et al., 2019). Indeed, we observed the noncanonical pathway expressed significantly elevated levels of intracellular RORγt at 72 hours compared to canonical treatment (**Fig. 3A**), supporting its involvement in the signaling cascade. To test the role of CBP/p300 in noncanonical T_H_17 polarization, we treated CD4+ T-lymphocytes polarized with canonical and noncanonical signals with A-485, a CBP/p300 inhibitor. IL-17A production was reduced by 88% in T-lymphocytes treated with the noncanonical signals and by 58% in T-lymphocytes treated with the canonical signals, compared to their respective controls (**Fig. 3B**). This suggests both pathways rely on CBP/p300 for IL-17A expression but may differ in the extent of that reliance.

**Figure 3:**
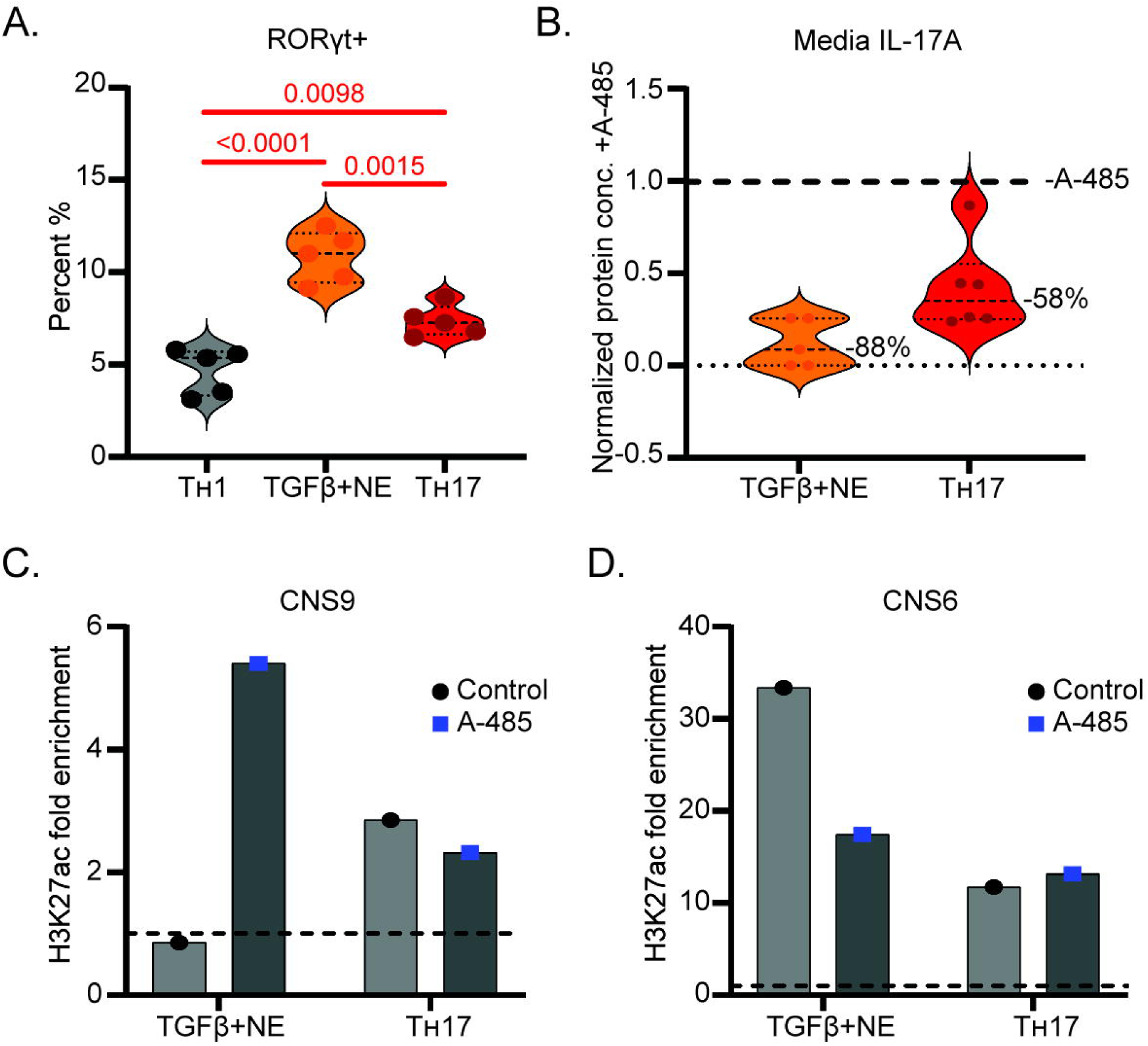
**CBP/p300 and ROR**γ**t are downstream targets of the noncanonical T_H_17 pathway.** Flow cytometry staining of intracellular RORγt 72 hours after T_H_1 (N=5), TGFβ+NE (N=5) and canonical T_H_17 (N=5) treatment of CD4+ T-lymphocytes **(A)**. Media IL-17A protein 72 hours after TGFβ+NE (N=5) and canonical T_H_17 (N=6) treatment of CD4+ T-lymphocytes treated with A-485, normalized to control activated CD4+ T-lymphocytes without A-485 (bold dotted line) **(B)**. Pooled (N=7) H3K27ac fold enrichment at the CNS9 and CNS6 loci normalized to naïve control T-lymphocytes (dotted line) **(C-D)**. Statistics were measured by Mann-Whitney (B) or one-way ANOVA with Tukey’s multiple comparisons tests (A).

The CBP/p300 complex is a primary driver of histone H3 acetylation at the H3K27 residue (Bedford and Brindle, 2012; Lasko et al., 2017; Tie et al., 2009), which is associated with open, activated chromatin at multiple regulatory elements of the Rorc locus, including the enhancers CNS6 and CNS9 (Chang et al., 2020; Friedman et al., 2023; Tian et al., 2021). CNS6 is a Rorc enhancer that is commonly associated with TGFβ/Smad signaling, while CNS9 is associated with IL-6-STAT3 signaling (Chang et al., 2020). To assess if these enhancers are involved in the noncanonical pathway, H3K27ac enrichment at the CNS6 and CNS9 loci was measured via chromatin immunoprecipitation. As expected, the canonical T_H_17 pathway exhibited higher H3K27ac enrichment at the CNS9 locus compared to the noncanonical pathway that lacks IL-6 signaling (**Fig. 3C**). However, the opposite was true for the CNS6 locus (**Fig. 3D**). Additionally, blocking CBP/p300 activity using A-485 blunted enrichment at the CNS6 locus but elevated enrichment at the CNS9 locus, while the canonical T_H_17 pathway was most unaffected. Together, these data further support the role of CBP/p300 in regulating RORγt in the noncanonical T_H_17 pathway, but a complex pattern of regulation has emerged that is in stark contrast to canonical signaling.

### RNA sequencing reveals distinct mechanisms between canonical and noncanonical T_H_17 pathways

Given the contrasting differences observed in these pathways, bulk RNA sequencing was performed at 24 hours post-treatment for the canonical and noncanonical T_H_17 pathways to obtain a more comprehensive analysis of their diverging mechanisms. Confirming our initial data, the two pathways varied greatly in transcriptional activity with tight clustering within the individual pathways, but significant variance across pathways as evidenced by PCA analysis (**Fig. 4A**). Unbiased clustering also demonstrated tight and reproducible gene expression distributions within the respective pathways and virtually opposite gene expression patterns across paired samples (**Fig. 4B**). Of note, Rorc, Batf, and Hif1α, which are known transcriptional regulators of canonical T_H_17 polarization (Dang et al., 2011; Ivanov et al., 2006; Schraml et al., 2009), are significantly elevated in the canonical T_H_17 compared to the noncanonical pathway at this timepoint (**Fig. 4B**), while approximately 1940 other genes are elevated in the noncanonical pathway that appear associated with GPCR signaling pathways (**Fig. 4C-D**). These data further suggest distinct programs between the pathways that lead to IL-17A expression, and lay the foundation for future investigation of the underlying mechanisms at hand.

**Figure 4:**
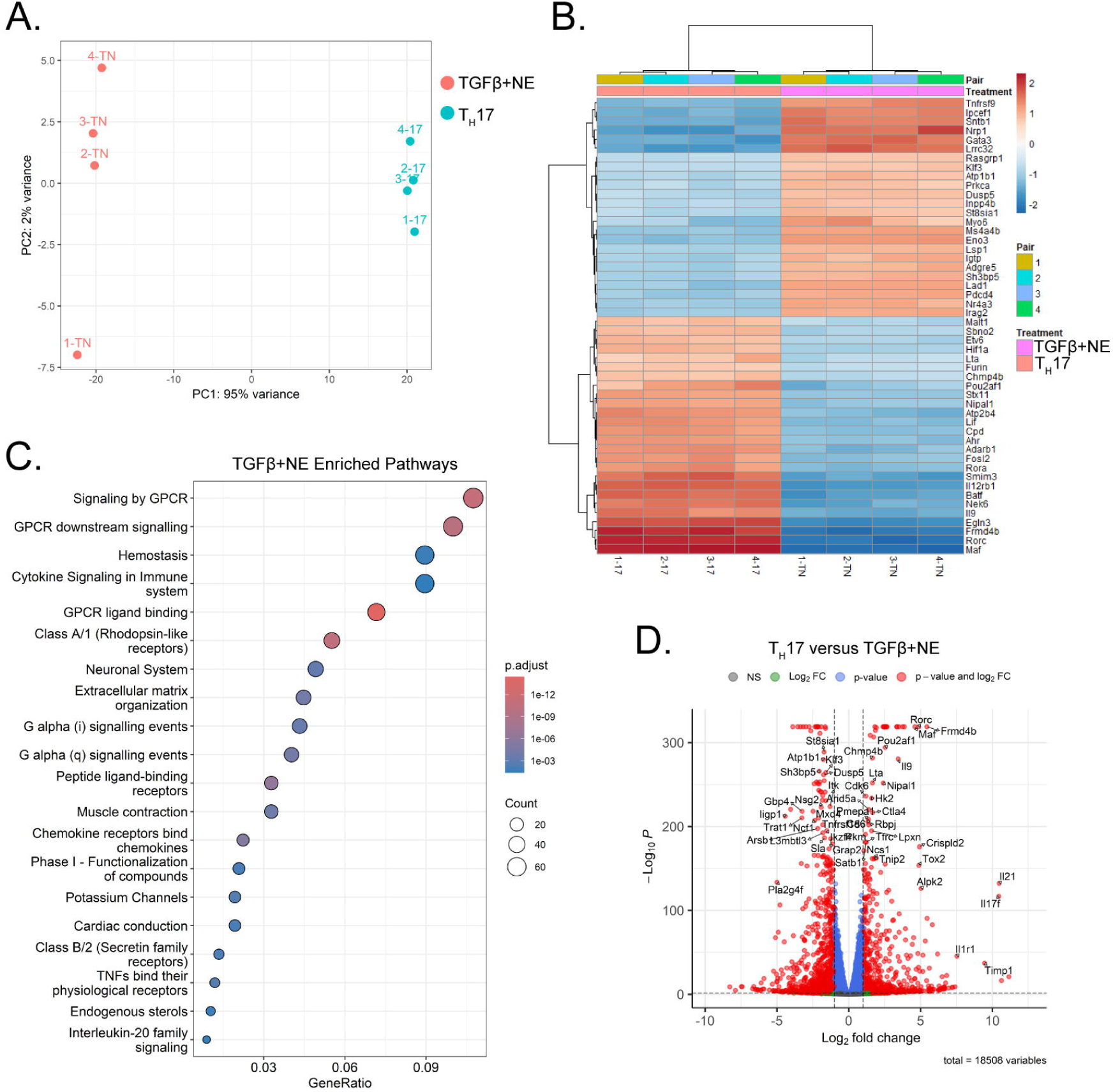
Bulk RNA sequencing data comparing canonical vs. noncanonical T_H_17 pathways. PCA analysis **(A)**, heat map of most significantly altered genes **(B)**, gene enrichment analysis **(C)**, and volcano plot **(D)** displaying the divergent genes and pathways involved in the canonical T_H_17 pathway and the TGFβ+NE pathway.

**Figure 5:**
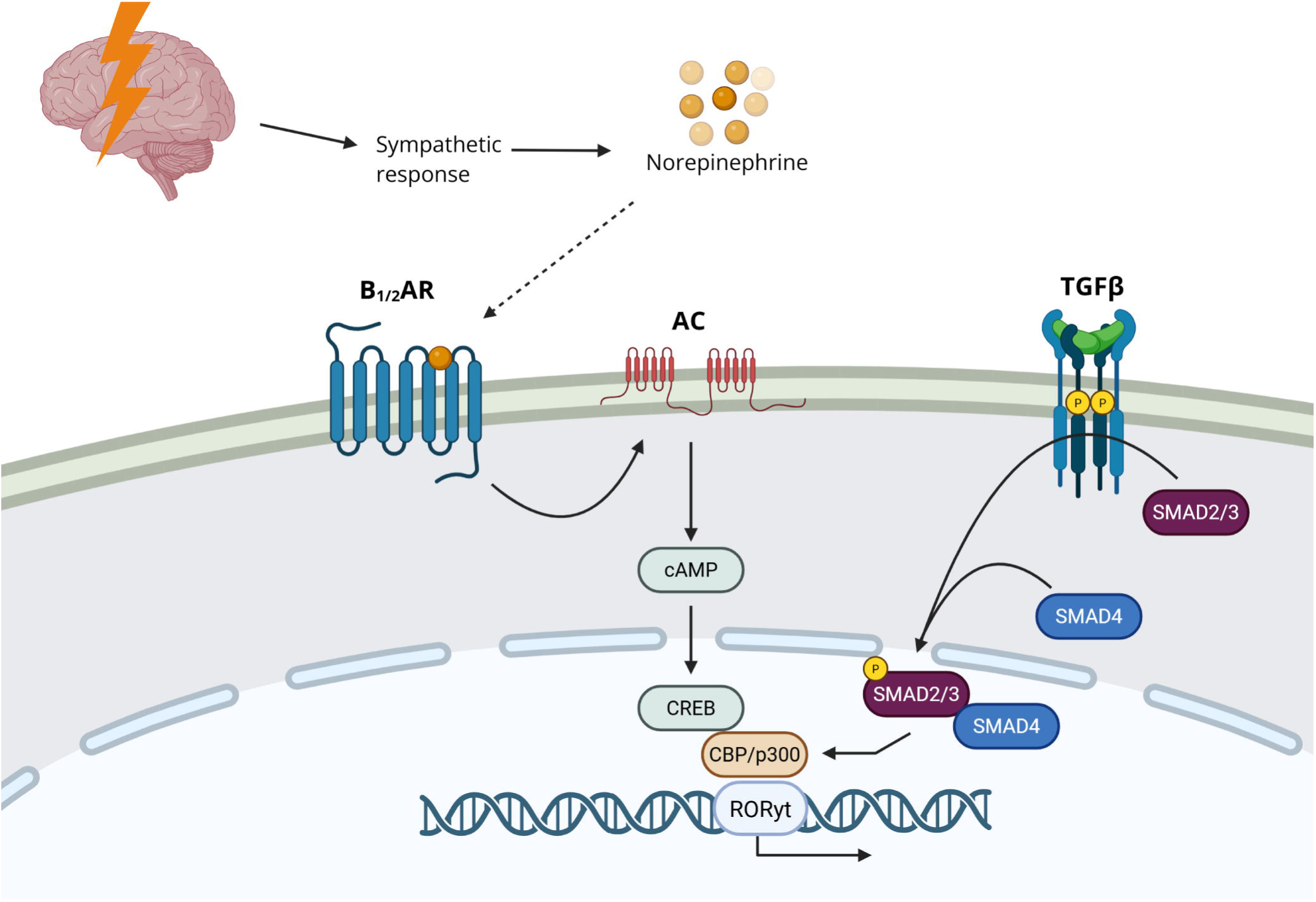
Proposed schematic representation of the noncanonical T_H_17 pathway. The sympathetic nervous system, signaling by NE, works through β-adrenergic receptors to signal via cAMP and CREB. CREB appears to transactivate CBP/p300 (an additional target of TGFβ) to activate the transcription of RORγt, a known regulator of IL-17A.

## Discussion

Our knowledge of the neuroimmune axis is constantly evolving. Currently, it is well-established that immune cells can both produce and respond to neurotransmitters, while neurons are able to produce and respond to cytokines (Nance and Sanders, 2007; Padro and Sanders, 2014). However, the role of neurotransmitters in T-lymphocyte polarization, particularly T_H_17, has not previously been reported. Our work presented herein has uncovered a novel role for NE in the regulation of T-lymphocyte IL-17A expression, which suggests that pathologies associated with dysregulated NE levels may be subject to concurrent altered systemic IL-17A levels.

Our work has previously demonstrated the necessity for β-adrenergic signaling and even T-lymphocyte-produced catecholamines for T-lymphocyte IL-17A expression (Elkhatib et al., 2022; Lauten et al., 2024), but the mechanisms underlying this signaling remained unclear. One of the most striking findings of the work presented herein is the necessity for endogenous peripheral catecholamines in the maintenance of systemic IL-17A levels. By reducing systemic catecholamine levels via 6-OHDA treatment, we were able to essentially eliminate detectable circulating levels of IL-17A. This suggests that basal sympathetic tone is intimately related to maintenance of IL-17A levels. Conversely, infusion of IL-17A has been shown to stimulate sympathetic nervous system (SNS) activity (Cao et al., 2021), implying the potential for a regulatory loop between the two neuroimmune entities. Furthermore, NE appears to be at the heart of regulating the relationship between IL-17A and the SNS. While 6-OHDA eliminates sympathetic nerve terminals that contain other catecholamines, including dopamine and epinephrine, it is important to note that NE is the primary and most abundant signaling neurotransmitter produced by the peripheral SNS (Klein Wolterink et al., 2022; Kostrzewa and Jacobowitz, 1974). Furthermore, the effects of 6-OHDA on epinephrine are minimal, as its primary source, the adrenal medulla, is largely unaffected by chemical sympathectomy (Caramona and Soares-da-Silva, 1985). Therefore, taken into aggregate, our data strongly supports the necessity of NE in IL-17A production.

Another critical finding of this work is that NE does not drive T-lymphocyte IL-17A alone, but requires concurrent TGFβ stimulation. While the primary *in vivo* source of NE is well-established to derive from the SNS, the source of TGFβ contributing to the noncanonical T_H_17 pathway is still unknown. A known source of circulating TGFβ is platelets (Assoian et al., 1983; Fava et al., 1990; Grainger et al., 1995; Meyer et al., 2012), which store TGFβ and release it upon activation. Indeed, pathologies like psychological stress have been reported to increase activated platelets in circulation (Andrén et al., 1983; Koudouovoh-Tripp and Sperner-Unterweger, 2012; Naesh et al., 1993), which may contribute the TGFβ necessary for the observed IL-17A increases in this disease. Macrophages are another possible source of TGFβ via β2-adrenergic signaling, which may be stimulated by stress-induced SNS activation (Yanagawa et al., 2016). However, we observed that basal IL-17A was regulated by NE, which based on our findings would also require TGFβ. Since both platelets and macrophages produce TGFβ in their activated or stimulated states, it remains unknown if these would be the sources for noncanonical T_H_17 polarization under homeostatic conditions. Given our disparate findings between canonical and noncanonical T_H_17 polarization, it could also suggest the source of TGFβ varies greatly between the two pathways as well. Additional work exploring the source of TGFβ in both health and disease may shed light on the mechanistic underpinnings of IL-17A regulation in context-specific conditions.

While this work suggests that TGFβ and NE together serve as regulators of homeostatic IL-17A production, it also suggests that dysregulation of these two factors (i.e., psychological stress or trauma) may also reset the IL-17A baseline, which could develop into a pathogenic T-lymphocyte response. A classic hallmark of this pathogenic T_H_17 phenotype resembles what has previously been termed the T_H_1/17 cell due to its ability to concurrently produce interferon-gamma (IFNγ) in addition to IL-17A (Harbour et al., 2015; Kamali et al., 2019; Tsanaktsi et al., 2018). While this T_H_1/17 subtype has previously been characterized by its cytokine production and involvement in disease, little is known about its source, lineage, differentiation, and function. Intriguingly, under high NE levels, our noncanonical T_H_17 pathway has repeatedly shown increases in IFNγ production (data not shown), unlike canonical T_H_17 polarization. Additionally, in our RNA sequencing analysis, STAT1 was significantly elevated in the noncanonical compared to the canonical T_H_17 pathway. STAT1 is a transcription factor known to drive T_H_1 polarization (Ai et al., 2022; Banik et al., 2021; Zhong et al., 2025) and is often suppressed by STAT3 in canonical T_H_17 polarization. This suggests that the noncanonical pathway that signals via TGFβ and NE may have the potential to become pathogenic under specific conditions, which may underlie the aforementioned association of chronic inflammatory conditions with numerous sympathoexcitation-associated psychopathologies.

In addition to STAT1, the involvement of STAT3 was also starkly different between the canonical and noncanonical pathways. As previously discussed, STAT3 was found not to be involved in the noncanonical T_H_17 pathway. This may be due to IL-6-driven STAT3 induction in the canonical pathway (Nishihara et al., 2007; Yang et al., 2007), a signal not present in the proposed pathway. Because STAT3 significantly and directly promotes RORγt expression (Durant et al., 2010; Yang et al., 2007; Yoon et al., 2015), we hypothesize that this is the rate-limiting difference between the canonical and noncanonical T_H_17 pathways – while the noncanonical pathway relies exclusively on the synergistic activity of CBP/p300 for RORγt transcription (and hence the larger response to A-485 inhibition), IL-17A production in the canonical pathway is driven both by TGFβ-mediated CBP/p300 activation, and more importantly, the robust STAT3 signal. Thus, the canonical T_H_17 pathway produces a much larger and faster IL-17A response due to its activation by rapidly produced cytokine milieus resulting from infections or direct foreign insults. In contrast, the noncanonical T_H_17 pathway appears to respond more slowly to allostatic changes such as sustained SNS activity. The temporal differences between the pathways are further shown by the elevated expression of RORγt in the canonical T_H_17 pathway at 24 hours (**Fig. 4B, D**), which seems to resolve by 72 hours, while the noncanonical pathway begins to express higher levels of RORγt at this later time (**Fig. 3A**).

This work has significant relevance for individuals with pathologies with elevated sympathoexcitation. For example, we have previously demonstrated the ability for β-blockade to completely prevent disease progression in experimental autoimmune encephalomyelitis (EAE), a mouse model of multiple sclerosis that is highly regulated by IL-17A (Lauten et al., 2025). In this work, we show that T-lymphocyte-specific β-blockade is sufficient to lessen IL-17A and EAE progression, which suggests that sympathoexcitation and NE contribute to the disease. Additionally, cardiovascular diseases, such as hypertension and heart failure, are known to possess elevated sympathoexcitation and IL-17A levels (Grassi et al., 2018; Liu et al., 2017). Interestingly, β-blockade is a primary mode of therapy in these diseases, which is primarily attributed to its direct impact on the heart. However, our findings here would assert that the lowering of IL-17A may also pose a therapeutic benefit, and our laboratory is currently investigating this hypothesis. Last, sympathoexcitation-associated psychopathologies, such as psychological trauma and PTSD, also have demonstrated increased levels of systemic IL-17A and elevated risks of inflammatory disorders. Interestingly, β-blockade using pharmaceuticals, such as propranolol, have been utilized in PTSD with little to no clinical benefit (C and R., 2020). However, in each of these clinical trials, propranolol’s effectiveness was made solely on the basis of behavioral changes. While this is logical given the specific disease being studied and the time these trials were performed (i.e., late 1990’s, early 2000’s – prior to the understanding that PTSD was linked to inflammation), no conclusions can be made as to the potential effectiveness of β-blockade on inflammation or risk of inflammatory disorders given the trial design. Moreover, we and others have recently observed and reported that the behavioral and pathophysiological sequalae of psychological trauma are not fully coupled (Case et al., 2025; Sumner et al., 2023). In other words, treatment of pathophysiology may not impact psychopathology, or vice versa. Given that the current standards of therapy for PTSD are treatments solely designed to ameliorate behavioral pathology, our findings here would suggest the need for additional studies (or revisiting) of adrenergic blockade as a potential therapeutic modality for the attenuation of psychological-trauma mediated inflammation.

In summary, we have uncovered a previously undescribed mechanism of T-lymphocyte IL-17A expression that utilizes NE-mediated neurotransmitter stimulation. This exciting discovery adds to our understanding of neuroimmune regulation and may provide a novel therapeutic target for IL-17A in an array of diseases.

## Acknowledgements and funding sources

This work was supported by the National Institutes of Health (NIH) R01HL158521 (AJC). We thank Texas A&M University’s Rodent Preclinical Phenotyping Core for the use of the Protein Simple Jess Automated Western Blot. We also thank Texas A&M University’s Institute for Genome Sciences and Society for bulk RNA sequencing and analysis.

## Author contributions

TN and AJC designed research studies. All authors conducted experiments, acquired data, and/or performed analyses. TN and AJC wrote the manuscript, while all authors approved the final version of the manuscript. AJC provided funding and experimental oversight.

## Abbreviations

RSDS: repeated social defeat stress
NE: norepinephrine
EAE: experimental autoimmune encephalomyelitis
IL6^-/-^: whole-body IL-6 knock out mice

---

**Supplementary Figure 1: NE and IL-17A are directly correlated.** Spleen **(A)** and plasma **(B)** NE depletion was confirmed via ELISA. Linear regression showing a positive correlation between plasma IL-17A and NE levels (N=4) **(C)**. Statistics were measured by Welch’s t-test (A,B) and linear regression (C).

