## Supplementary figures and images for "Norepinephrine is a novel and essential regulator for T-lymphocyte interleukin 17A expression"

### Supplemental Figure 1

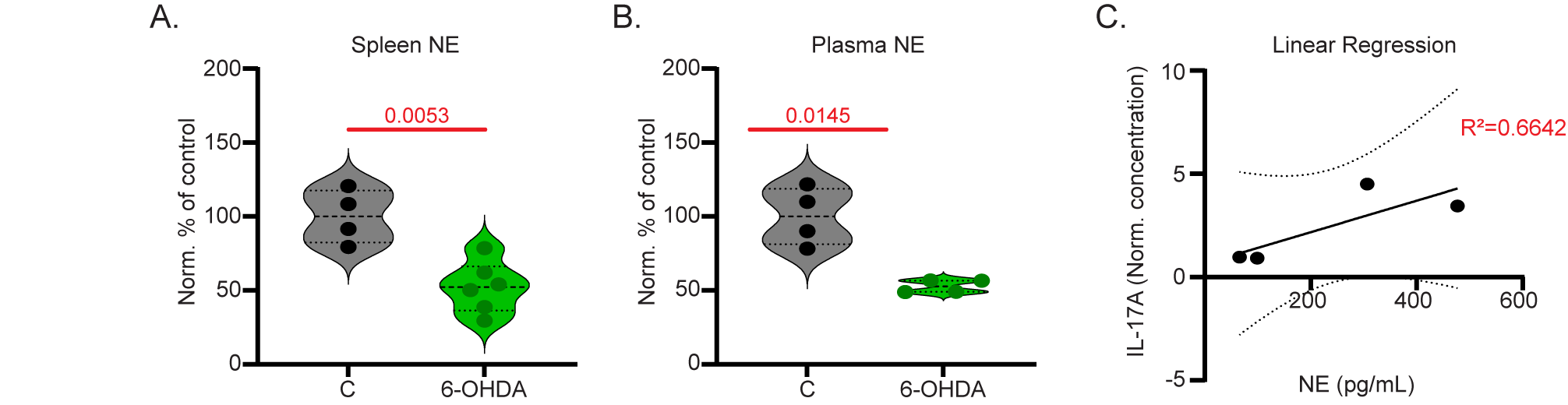
